# A Novel Heme-Degrading Enzyme that Regulates Heme and Iron Homeostasis and Promotes Virulence in *Enterococcus faecalis*

**DOI:** 10.1101/2025.01.20.633879

**Authors:** Debra N. Brunson, Hader Manzer, Alexander B. Smith, Joseph P. Zackular, Todd Kitten, José A. Lemos

## Abstract

*Enterococcus faecalis*, a gut commensal, is a leading cause of opportunistic infections. Its virulence is linked to its ability to thrive in hostile environments, which includes host-imposed metal starvation. We recently showed that *E. faecalis* evades iron starvation using five dedicated transporters that collectively scavenge iron from host tissues and other iron-deprived conditions. Interestingly, heme, the most abundant source of iron in the human body, supported growth of a strain lacking all five iron transporters (Δ5Fe). To release iron from heme, many bacterial pathogens utilize heme oxygenase enzymes to degrade the porphyrin that coordinates the iron ion of heme. Although *E. faecalis* lacks these enzymes, bioinformatics revealed a potential ortholog of the anaerobic heme-degrading enzyme anaerobilin synthase, found in *Escherichia coli* and a few other Gram-negative bacteria. Here, we demonstrated that deletion of OG1RF_RS05575 in *E. faecalis* (ΔRS05575) or in the Δ5Fe background (Δ5FeΔRS05575) led to intracellular heme accumulation and hypersensitivity under anaerobic conditions, suggesting *RS05575* encodes an anaerobilin synthase, the first of its kind described in Gram-positive bacteria. Additionally, deletion of *RS05575*, either alone or in the Δ5Fe background, impaired *E. faecalis* colonization in the mouse gastrointestinal tract and virulence in mouse peritonitis and rabbit infective endocarditis models. These results reveal that RS05575 is responsible for anaerobic degradation of heme and identify this relatively new enzyme class as a novel factor in bacterial pathogenesis. Findings from this study are likely to have broad implications, as homologues of *RS05575* are found in other Gram-positive facultative anaerobes.

**IMPORTANCE:** Heme is an important nutrient for bacterial pathogens, mainly for its ability to serve as an iron source during infection. While bacteria are known to release iron from heme using enzymes called heme oxygenases, a new family of anaerobic heme-degrading enzymes has been described recently in Gram-negative bacteria. Here, we report the first description of anaerobic heme degradation by a Gram-positive bacterium, the opportunistic pathogen *Enterococcus faecalis*, and link activity of this enzyme to their ability to colonize and infect the host. We also show that homologues of this enzyme are found in many Gram-positive facultative anaerobes, implying that the ability to degrade heme under anaerobic conditions may be an overlooked fitness and virulence factor of bacterial pathogens.

## INTRODUCTION

*Enterococcus faecalis* is a facultative anaerobe known for its intrinsic multi-stress resiliency and ability to cause numerous opportunistic infections (1). However, *E. faecalis* is also a member of the gut microbiota and will typically only cause disease under specific conditions that include, but are not limited to, extended antibiotic usage, compromised immunity, and utilization of indwelling medical devices (2). For the most part, the virulence of *E. faecalis* derives from its inherent capacity to overcome various stress conditions, form biofilms on both biotic and abiotic surfaces, and subvert immune responses (3). Regarding the latter, a central aspect of the innate immune response of enterococcal hosts involves the rapid mobilization of proteinaceous metal chelators to the site of infection that avidly bind to trace metals such as iron, manganese and zinc, a process known as nutritional immunity (4–7).

An essential trace metal to virtually all forms of life, iron holds a prominent role in bacterial physiology and in host-pathogen interactions as its electrochemical properties and abundance in nature makes it the preferred redox cofactor for enzymatic reactions (8). We recently showed that *E. faecalis* can efficiently scavenge iron from the environment via the cooperative activity of three highly conserved and two novel iron transporters (9). The simultaneous inactivation of all five transporters (Δ5Fe strain) resulted in major growth impairment under iron-depleted conditions, which, as expected, was accompanied by a substantial reduction in intracellular iron pools. However, the virulence potential of the Δ5Fe strain in animal models varied depending on the type of model (the invertebrate *Galleria mellonella* larvae or mice) and, in the case of mice, the infected site. Specifically, virulence of Δ5Fe was significantly attenuated in *G. mellonella*; however, the Δ5Fe strain showed impaired capacity to infect the peritoneal cavity while it disseminated and infected spleens as well as the parental strain. We suspected that these differences correlate with heme availability, as heme— plentiful in blood and mammalian tissues—serves as a major source of iron for some of the most successful bacterial pathogens (10–13). Furthermore, non-hematophagous insects like *G. mellonella* are virtually heme-free, as they use two copper ions coordinated by histidine residues to transport oxygen rather than the hemoglobin/Fe-heme complexes found in vertebrates (14).

As anticipated, heme supplementation restored growth and intracellular iron homeostasis in the Δ5Fe strain grown in media lacking any other type of iron source while injection of small amounts of heme into the *G. mellonella* hemocoel restored virulence of Δ5Fe to levels comparable to those of the parent strain (9). While these results clearly demonstrate that *E. faecalis* can utilize heme as an iron source, the mechanisms by which it acquires heme from the environment—since enterococci cannot synthesize heme (15, 16)—and how the iron ion is released from the porphyrin ring remain unknown.

Oxidative degradation mediated by heme oxygenases is the best described mechanism of heme degradation in bacteria (17). In important and diverse bacterial pathogens such as *Streptococcus pyogenes* and *Pseudomonas aeruginosa*, the canonical HO-1 heme oxygenase uses oxygen to disrupt the tetrapyrrole ring to liberate biliverdin, CO_2_ and Fe^+2^ (17, 18).

Additionally, some bacteria encode the so-called non-canonical heme oxygenase, such as the *Staphylococcus aureus* IsdG/I, which uses oxygen to degrade heme into staphylobilin and formaldehyde (19–21). Recently, an oxygen-sensitive radical S-adenosylmethionine methyltransferase (rSAM), named ChuW, was identified in *Escherichia coli*, and shown to degrade heme’s porphyrin ring under anaerobic conditions (22). ChuW utilizes a primary carbon radical S-adenosylmethionine to promote methyl transfer and subsequent linearization of the porphyrin ring, releasing the iron atom and the linear tetrapyrrole product anaerobilin, hence the designation anaerobilin synthase (22, 23). Aside from *E. coli*, ChuW homologues have been described in *Vibrio cholerae* and *Fusobacterium nucleatum* (22, 24, 25). Considering that either canonical or non-canonical heme oxygenases cannot function under anaerobic conditions, the presence of an oxygen-independent enzyme that mediates heme degradation is expected to provide a competitive advantage for bacteria that inhabit anaerobic environments.

Through bioinformatic analysis, we identified a ChuW ortholog in the *E. faecalis* OG1RF genome. Further analysis indicated that OG1RF_RS05575 (hereafter referred to as RS05575) is conserved across the *Enterococcus* genus as well as other facultative anaerobic Gram-positive cocci (26). In this investigation, we provide the first insights into the mechanisms of anaerobic heme degradation in Gram-positive bacteria and, for the first time, link the activity of an anaerobilin synthase with bacterial virulence.

## RESULTS

### *Enterococcus faecalis* internalizes and utilizes heme as an iron source

We recently showed that *E. faecalis* utilizes heme as an iron source while others have shown that it will grow to higher cell density and form more robust biofilms in the presence of heme (9, 27). To further explore the relationship between heme and iron in *E. faecalis*, we tested whether heme supplementation could hinder elemental iron uptake. To do so, *E. faecalis* OG1RF was grown to mid-log phase in a chemically-defined medium, FMC, lacking an iron source (28). Then, mid-log grown cultures were divided into aliquots and supplemented with either 1µM ^55^Fe (control), 1µM ^55^Fe + 10µM heme, or 1µM ^55^Fe + 40µM FeSO_4_ with samples taken after 1 and 5 minutes. As expected, the addition of cold (unlabeled) FeSO_4_ effectively slowed ^55^Fe uptake with ∼70% reduction after 5 minutes when compared to the control sample (Fig 1A). Heme was also effective, seemingly more than FeSO_4_, as it reduced ^55^Fe uptake by ∼90% after the same period (Fig 1A). As we have shown that transcription of the iron transporters *efaABC*, *emtABC*, *feoAB, fhuDCBG* and *fitABCD* was significantly elevated during iron starvation (9), we next asked if heme supplementation could shut down this activation. Indeed, heme treatment lowered transcription of *feoB* (∼3-fold), *fitA* (∼15-fold)*, fhuB* (∼13-fold), and *emtB* (∼1.5-fold) (Fig 1B). However, heme treatment led to ∼100-fold increase in *efaA* expression (Fig 1B). Because *efaCBA* codes for a dual iron/manganese transporter, we wondered if this induction was necessary to enhance manganese uptake to mitigate heme toxicity and assure maintenance of a balanced iron:manganese ratio. To investigate this possibility, we assessed *mntH2* levels, the other major manganese transporter of *E. faecalis* (29) and of the heme efflux pump *hrtA* (30). As predicted, *mntH2* levels were 10-fold higher after heme treatment while *hrtA* was induced by approximately 100-fold (Fig 1B). Finally, we determined intracellular heme content of OG1RF cultures grown in FMC supplemented with 20µM heme +/- 10µM FeSO_4_. We found that when grown with both heme and FeSO_4_, intracellular heme levels nearly doubled (∼73% increase) compared to cells grown only in heme indicating that *E. faecalis* will slow down heme degradation when free iron is available (Fig 1C). Collectively, these results provide unequivocal evidence that *E. faecalis* can rapidly import and then degrade heme to release the iron ion.

**Fig 1.**
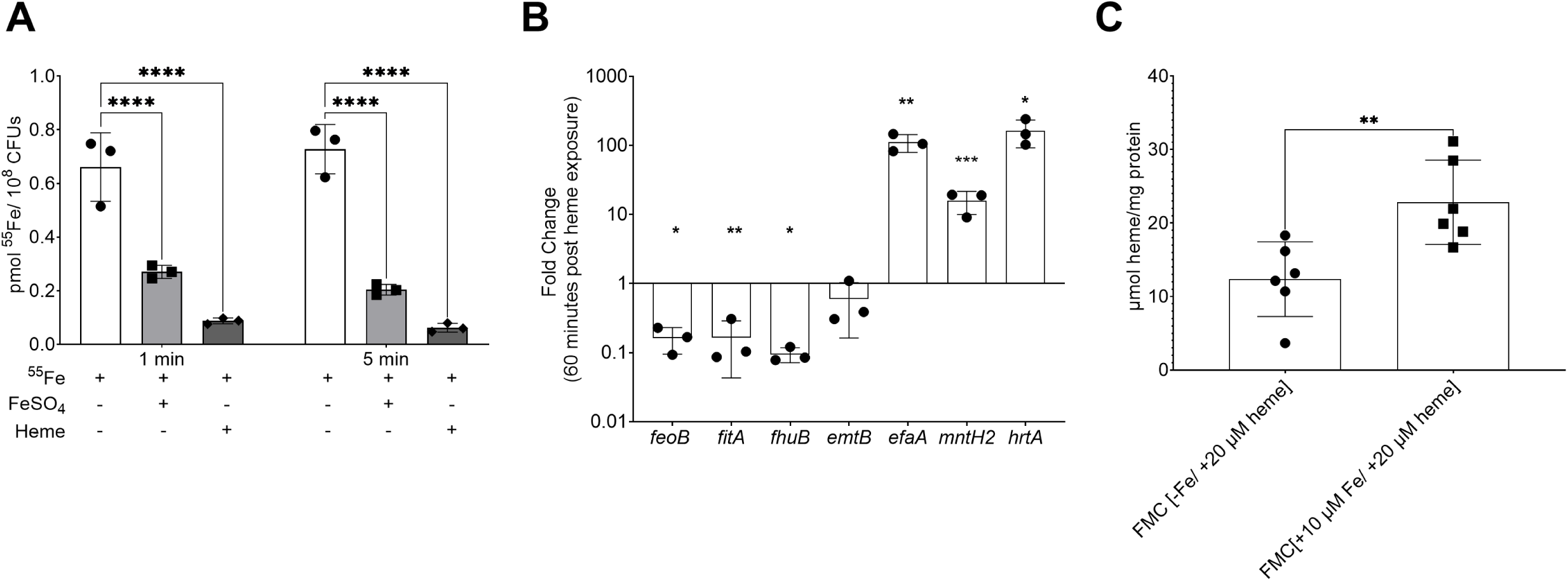
*Enterococcus faecalis* uses heme as an iron source. (**A**) ^55^Fe uptake by *E. faecalis* is diminished by competition with unlabeled heme and iron. OG1RF was grown in FMC[-Fe] to OD_600_ 0.5 and ^55^Fe uptake monitored at one and five minutes after addition of 1µM ^55^Fe, 1µM ^55^Fe + 10µM heme, or 1µM ^55^Fe + 40µM FeSO_4_. Statistical significance was determined by Two-Way ANOVA with Dunnett’s multiple comparison test. (**B**) RT-PCR analysis showing that heme supplementation represses iron uptake genes but activates transcription of manganese uptake genes. OG1RF was grown to OD_600_ 0.5 in FMC[-Fe] and sampled before and 60 minutes after supplementation with 20 µM heme. Statistical significance was determined by Student’s T- test. (**C**) Excess iron leads to increased intracellular heme levels. OG1RF was grown in either FMC[-Fe +20 µM heme] or FMC[+10 µM Fe +20 µM heme] to OD_600_ 0.5 and the intracellular heme content determined. Statistical significance was determined by a Student’s T-test, *p≤0.05, **p≤0.01, ****p≤0.0001.

### Identification of an oxygen-sensitive heme-degrading enzyme in *E. faecalis*

To serve as an iron source, the porphyrin ring of heme must be degraded. In bacteria, oxidative degradation is often mediated by enzymes called heme oxygenase. *In silico* analysis indicate that enterococcal genomes do not encode heme oxygenases (26) but, on the other hand, identified an ortholog of *E. coli* ChuW (gene ID: OG1RF_RS05575). ChuW is an oxygen-sensitive, r-SAM- type enzyme, that catalyzes the degradation of heme into a linear tetrapyrrole named anaerobilin (23). Even though the similarity between ChuW and RS05575 appears unremarkable (23% identity and 44.5% similarity), the CXXXCXXC r-SAM motif essential for binding of S-adenosylmethionine and an aspartic acid residue critical for ChuW activity (23) are conserved in RS05575, with several other important residues identified in ChuW and few other anaerobilin synthases characterized to date being also present in RS05575 (Fig 2B). Despite this moderate similarity at the amino acid level, superimposition of AlphaFold2 predicted structures of *E. coli* ChuW and *E. faecalis* RS05575 revealed a striking structural similarity between the two proteins (Fig 2C). Using a pairwise structure alignment tool from the Research Collaboratory for Structural Bioinformatics (RCSB) Protein Data Bank (31), we derived analytical scores to demonstrate the structural similarity of ChuW and RS05575 with a root mean square deviation score of 3.01 and a template modeling score of 0.78. Using the AlphaFold server and the HeMoQuest webserver to predict potential ligand interactions, we found that RS05575 likely binds heme using Y7, C20, Y59, Y190, Y236, and/or H250 as coordinating residues (Fig S1).

**Fig 2.**
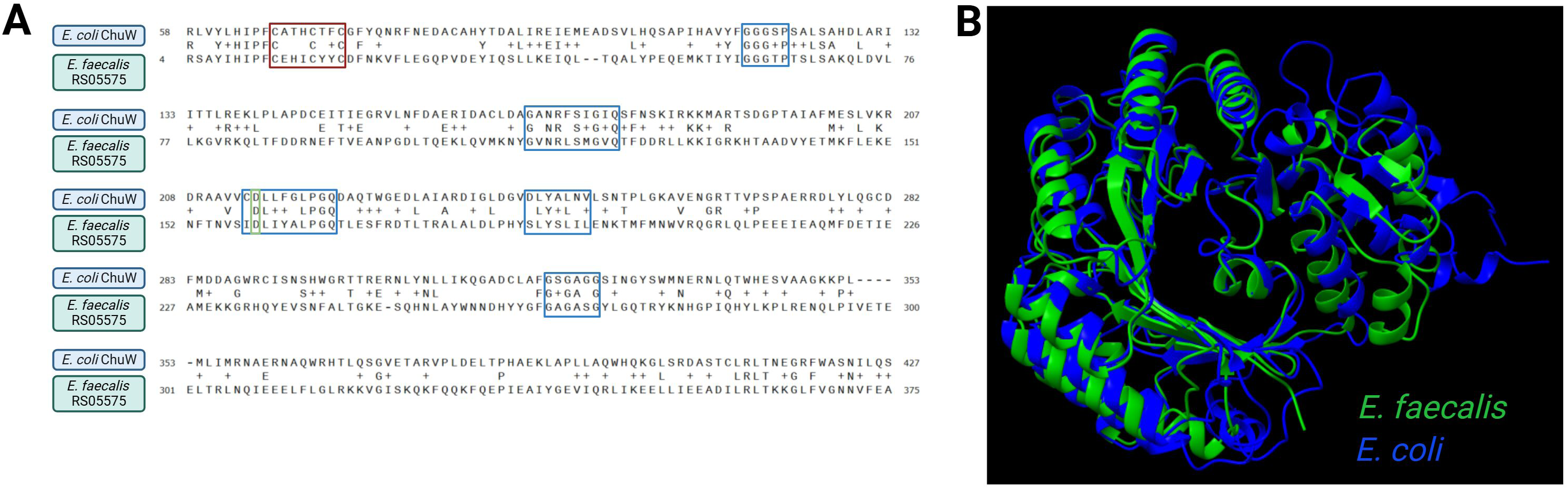
Identification of a putative anaerobilin synthase in *E. faecalis*. (**A**) Amino acid alignment of *E. faecalis* RS05575 and *E. coli* ChuW. The canonical rSAM CXXXCXXC motif is outlined in red, the aspartic acid residue known to be important for ChuW activity is outlined in green, and residues conserved across *E. coli* ChuW, *Vibrio cholerae* HutW, and *Fusobacterium nucleatum* HmuW are outlined in blue. (**B**) AlphaFold 2 structures of *E. faecalis* RS05575 and *E. coli* ChuW superimposed on each other using ChimeraX.

Analysis of all genome sequences available at the Bacterial and Viral Bioinformatic Resource Center (BV-BRC) (26) indicated that RS05575 is widespread among enterococci and members of the Enterococcaceae family (Fig 3 and Table S1). Notably, RS05575 orthologs with ∼70% similarities are found in several facultative Gram-positive anaerobes, including streptococcal and staphylococcal species.

**Fig 3.**
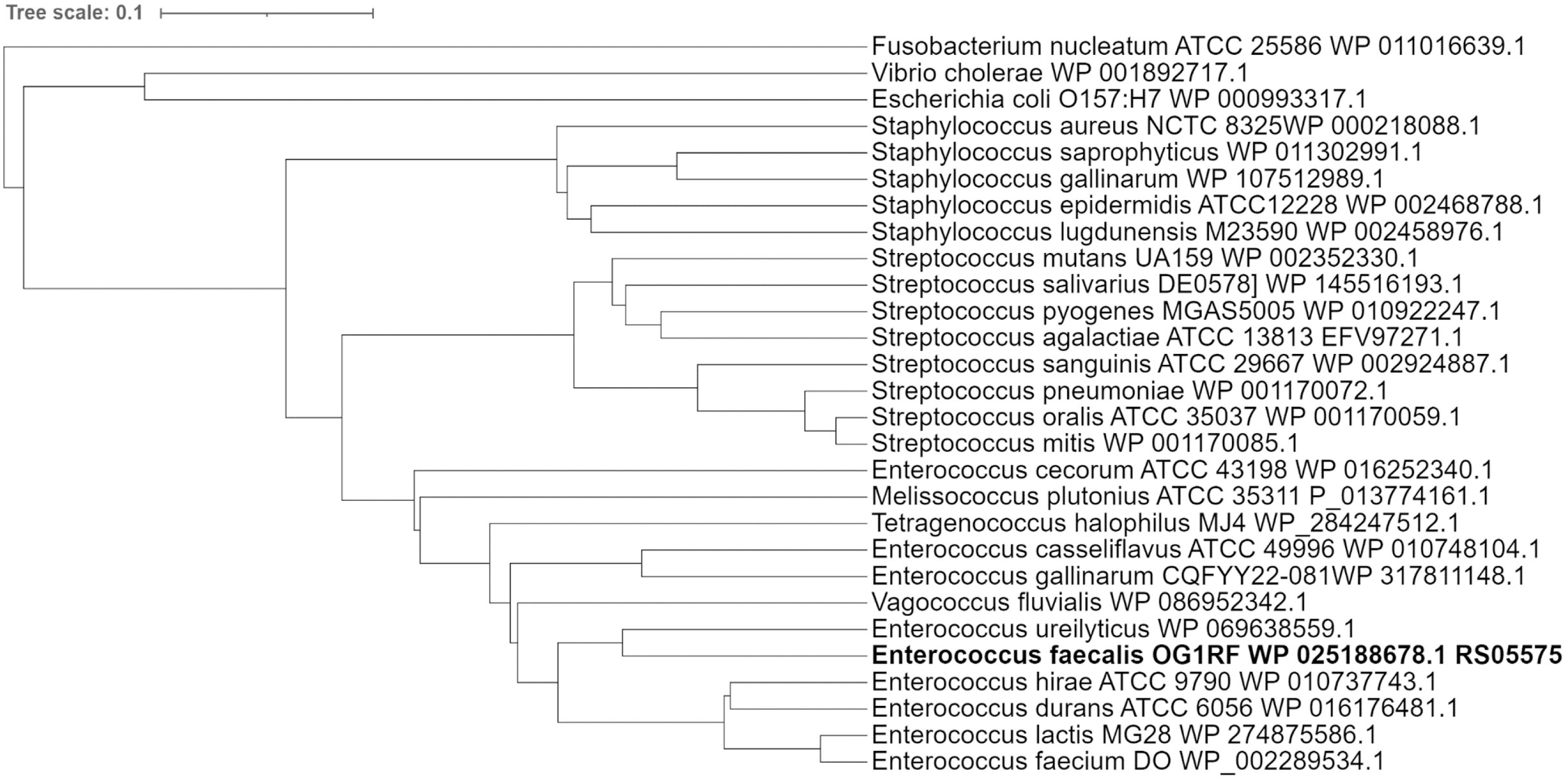
Phylogenetic analysis of RS05575 homologues in other Gram-positive facultative anaerobes and select Gram-negatives. BlastP searches against RS05575 were used to identify homologues across species of enterococci, streptococci, staphylococci, *E. coli*, *V. cholerae*, and *F. nucleatum*. Phylogenetic trees were constructed using multiple sequence alignments of representative species using Clustal Omega and iTOL.

### Inactivation of RS05575 differentially affects growth of *E. faecalis* under aerobic and anaerobic conditions

To explore the biological significance of RS05575, we generated a RS05575 deletion (ΔRS05575) and genetically complemented (ΔRS05575c) strains. We assessed growth of ΔRS05575 in FMC medium originally prepared without an iron source (FMC[-Fe]) and then supplemented with either 20 µM heme (FMC[+heme]) or 20 µM FeSO_4_ (FMC[+Fe]). As we expected RS05575 to be only active under anaerobiosis, we assessed the capacity of ΔRS05575 to grow under both aerobic and anaerobic conditions. For the latter, we used FMC retaining dissolved oxygen (herein FMC_DO_) as well as O_2_-purged media (FMC_O2P_). In FMC[-Fe], the ΔRS05575 strain grew as well as the parent, reaching slightly higher growth yields under the more strict (FMC_O2P_) anaerobic condition (Fig 4A-C). Similar results were obtained in FMC[+Fe] as the mutant reached higher final growth yields under anaerobiosis (Fig 4D-F). In FMC[+heme], ΔRS05575 strain grew similarly to the parent strain reaching higher growth yields when incubated in air but not in FMC_DO_ (Fig 4G-H). Notably, both the OG1RF parent and ΔRS05575 strains struggled to grow in the presence of heme under strict (FMC_O2P_) anaerobic conditions. Specifically, OG1RF grew very poorly with an extended lag phase of ∼12 hours while the ΔRS05575 strain failed to grow (Fig 4I). Genetic complementation restored all relevant growth phenotypes to parental levels (Fig 4C, G and I) of ΔRS05575.

**Fig 4.**
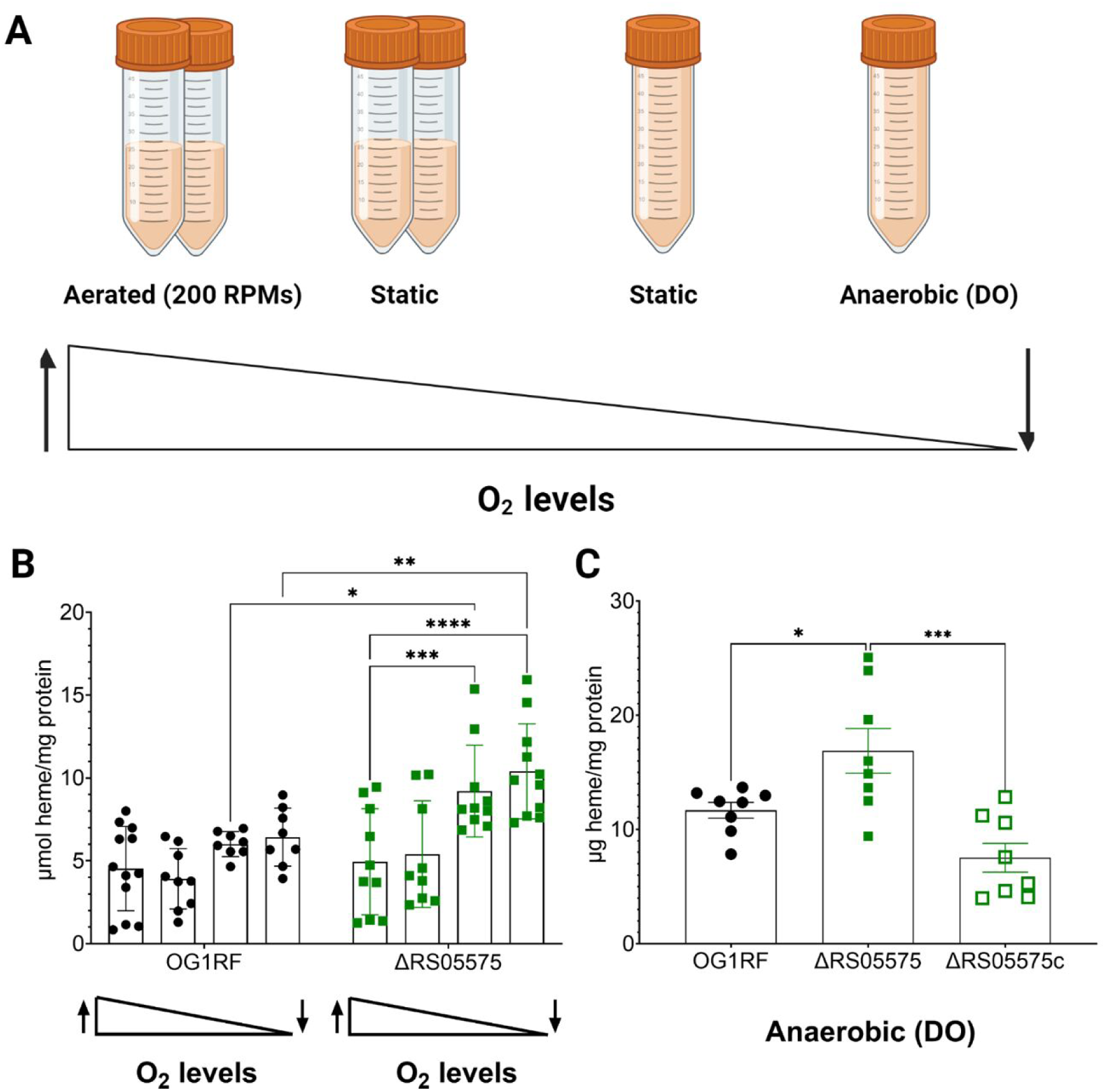
Growth of *E. faecalis* OG1RF and ΔRS05575 strains in media containing different amounts of FeSO_4_ and heme and under different atmospheres. (**A**) FMC[-Fe], (**B**) FMC[+20 µM Fe], (**C**) FMC[+20 µM heme], (**D**) FMC_DO_[-Fe], (**E**) FMC_DO_[+20 µM Fe], (**F**) FMC_DO_[+20 µM heme], (**G**) FMC_O2P_[-Fe], (**H**) FMC_O2P_[+20 µM Fe], and (**I**) FMC_O2P_[+20 µM heme]. Cells were grown overnight in FMC[-Fe], FMC_DO_[-Fe], or FMC_O2P_[-Fe], normalized to OD_600_ 0.2 and sub- cultured at 1:200 into the designated media. Growth was monitored by measuring OD_600_ every 30 minutes using an automated growth reader. The ΔRS05575c strain was used to show genetic complementation of the more noticeable phenotypes (**C**, **G** and **I**). Error bars denote standard error of the mean from at least two independent experiments with three biological replicates each.

### Inactivation of RS05575 leads to heme accumulation in microaerophilic and anaerobic environments

To investigate the role of RS05575 in heme catabolism, we quantified intracellular heme in the OG1RF and ΔRS05575 strains grown to mid-log phase in FMC[-Fe] supplemented with 20 µM heme under an oxygen gradient (see methods and Fig 5A for details) While dissolved oxygen levels had no impact on heme pools in OG1RF, intracellular heme nearly doubled in ΔRS05575 grown under low oxygen (static with no headspace) or anaerobic conditions (Fig 5B). Genetic complementation reversed this phenotype (Fig 5C).

**Fig 5.**
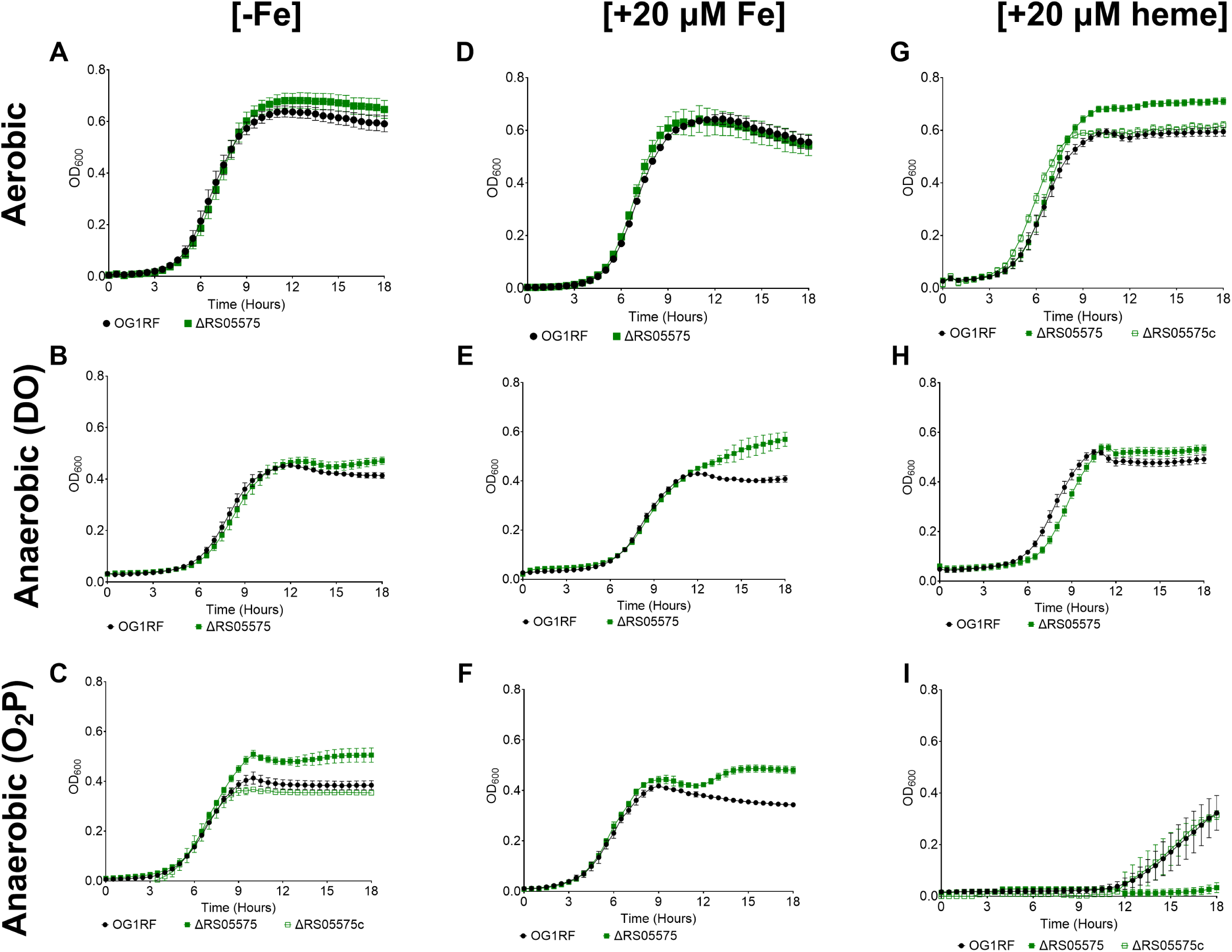
RS05575 degrades heme under oxygen-depleted conditions. (**A**) Strains were grown in FMC[+20 µM heme] to OD_600_ ∼0.5 in a shaking incubator with 50% headspace, a static incubator with 50% headspace, a static incubator with no headspace, or in the anaerobic chamber with no headspace. All media contained dissolved oxygen to bypass the extreme growth defect under anaerobic conditions in the presence of heme. (**B, C**) Intracellular heme content of OG1RF, ΔRS05575, and ΔRS05575c. Individual biological replicates from at least 2 independent experiments shown with n≥8. Error bars denote standard error of the mean. Statistical significance was determined by a Two-way ANOVA with Šidák’s multiple comparison test,*p≤0.05, **p≤0.01, ***p≤0.001 ****p≤0.0001.

To further define the role of RS05575 in heme degradation and its importance to iron homeostasis, we leveraged the extreme iron starvation that can be imposed to the Δ5Fe strain by growing cells in iron-depleted media (9). Specifically, we generated a sextuple mutant by introducing the RS05575 deletion into the Δ5Fe background and used the original Δ5Fe as well as the ΔRS05575 and Δ5FeΔRS05575 strains to compare their heme uptake and degradation efficiency. When compared to Δ5Fe, the Δ5FeΔRS05575 strain grew equally well in different oxygen content under iron-starving conditions or in media supplemented with either FeSO_4_ or heme (Fig. S1A-I). However, different than ΔRS05575, the Δ5FeΔRS05575 strain grew in FMC_O2P_[+heme] albeit still displaying an extended lag phase. The reason for this unexpected observation remains to be determined.

Upon identifying conditions that supported growth of all strains, we next monitored their heme uptake capacity by growing cells to mid-log phase in FMC_O2P_[-Fe], spiked cultures with 10µM heme, and monitored intracellular heme content 5, 15, and 60 minutes after heme treatment. As expected based on prior evidence that the Δ5Fe strain is primed to take up heme (9), both Δ5Fe and Δ5FeΔRS05575 acquired heme much more rapidly than the OG1RF and ΔRS05575 strains (Fig 6A, notable differences at T_15-min_). To monitor heme degradation, we set up another experiment where cultures were spiked with 10 µM heme for 15 minutes, the cells collected by centrifugation, washed in PBS once, and suspended in fresh FMC_O2P_[-Fe] with intracellular heme monitored for up to 3 hours. While intracellular heme continued to increase in both OG1RF and ΔRS05575 during the first hour after media change, likely due to residual uptake of heme bound to the cell surface, it declined by ∼35% in OG1RF after 3 hours while remaining steady in ΔRS05575 (Fig 6B). Most importantly, heme levels sharply decreased (∼65%) in the Δ5Fe strain after 3 hours but not in Δ5FeΔRS05575. Collectively, these results strongly support that RS05575 mediates anaerobic heme degradation (Fig 6B).

**Fig 6.**
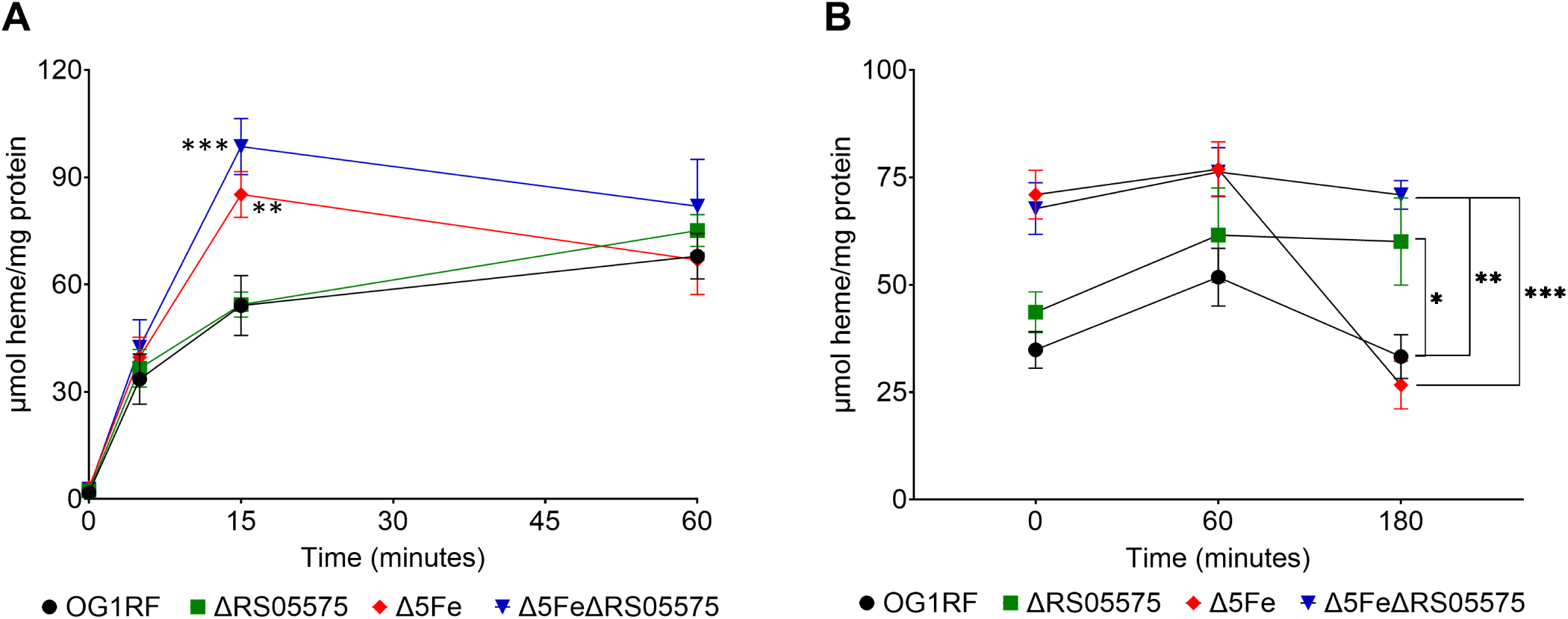
Uptake and degradation of heme by RS05575 in anaerobic conditions. (**A**) OG1RF, ΔRS05575, Δ5Fe, and Δ5FeΔRS05575 were grown to OD_600_ ∼0.5 in FMC_O2P_[-Fe] at which point cultures were supplemented with 10 µM heme and the intracellular heme determined after 0, 5, 15, and 60 minutes. (**B**) Cultures were supplemented with 10 µM heme for 15 minutes, washed, and cell pellets suspended in FMC_O2P_[-Fe]. Samples were taken at 0, 60, and 180 minutes after heme removal. All data was normalized to protein content. Experiments were performed with 3 biological replicates on at least two independent occasions. Error bars denote standard error of the mean. Statistical significance was determined by an ordinary One-way ANOVA with Dunnett’s multiple comparisons test at each time point,*p≤0.05, **p≤0.01, ***p≤0.001 ****p≤0.0001.

### RS05575 plays a role in *E. faecalis* virulence and intestinal colonization

Upon demonstration that RS05575 mediates heme degradation under anaerobic conditions, we sought to investigate its possible role in enterococcal fitness and pathogenesis. Given the close association between heme and iron homeostasis, we conducted the following series of experiments using the ΔRS05575, Δ5Fe, and Δ5FeΔRS05575 strains. First, we used an intra- peritoneal challenge mouse model, in which *E. faecalis* spreads systemically within 24 hours. As shown previously (9), the ability of the Δ5Fe strain to infect the peritoneal cavity was impaired (∼1-log reduction) when compared to OG1RF but not in the (heme-rich) spleen (Fig 7A-B). We also showed that the Δ5Fe strain could efficiently colonize the heart and liver but not the kidney (Fig 7C-E). The ΔRS05575 single mutant displayed defective ability to infect the peritoneal cavity, liver and kidney, but not spleen or heart. Finally, virulence of Δ5FeΔRS05575 was attenuated in all tissues sampled and was the only mutant recovered at significantly lower numbers from spleens and hearts when compared to the OG1RF parent strain (Fig 7).

**Fig 7.**
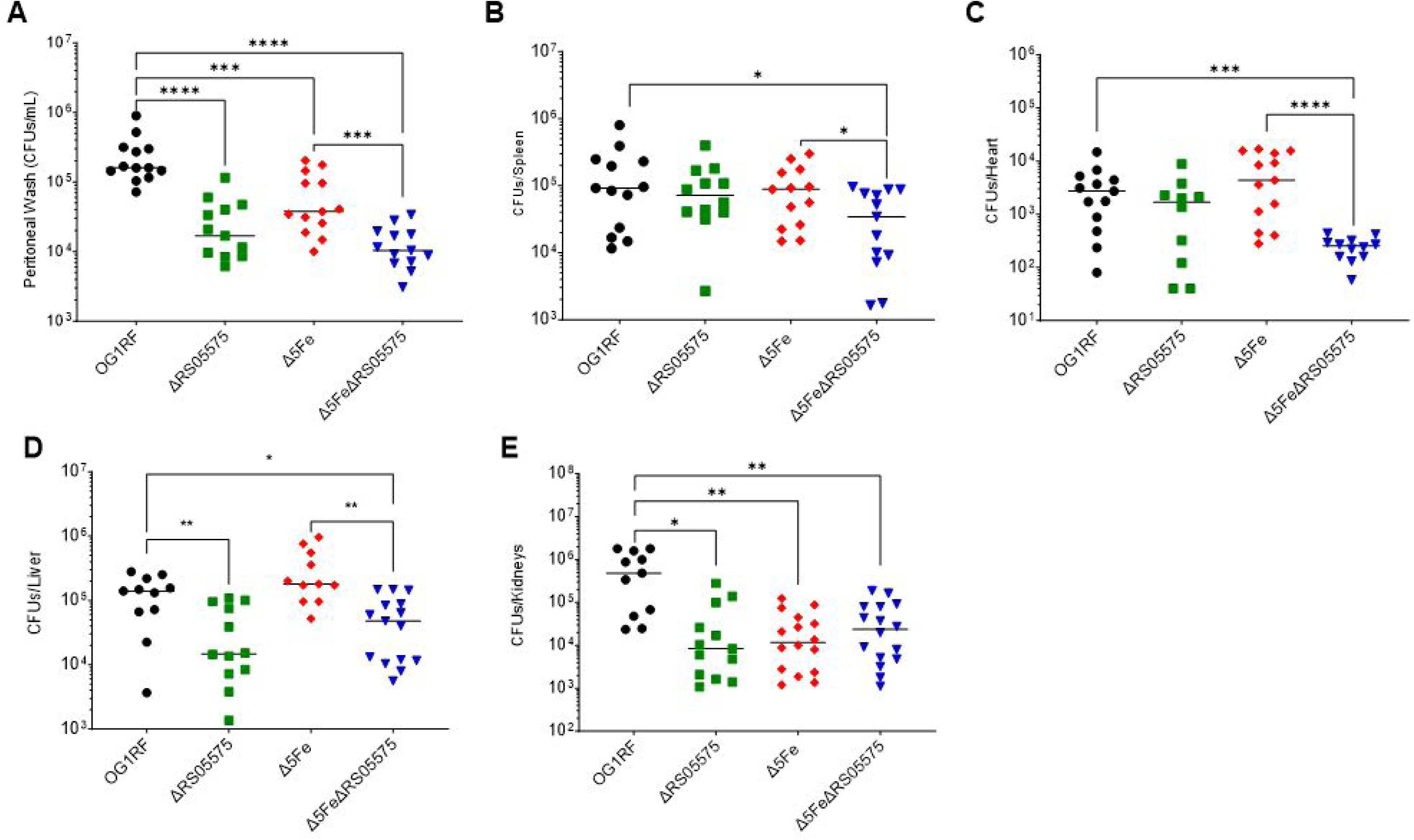
Loss of RS05575 and all 5 iron transporters (Δ5Fe strain) alone or in combination differentially affects *E. faecalis* virulence potential. 7-week-old C57BL6-J mice from Jackson Laboratories were infected with 1X10^8^ CFUs via intraperitoneal injection and the mice euthanizes after 48 hours and the different tissues collected for CFU determination. (**A**) peritoneal wash, (**B**) spleens, (**C**) hearts, (**D**) livers, and (**E**) kidneys. The data points shown are a result of the ROUT outlier test and bars denote median values. Statistical analyses were performed using the Mann-Whitney test, *p≤0.05, **p≤0.01, ***p p≤0.001, and ****p≤0.0001.

Next, we used the rabbit infective endocarditis (IE) model to determine if RS05575 also plays a role in enterococcal IE and, in parallel, assess the virulence of the Δ5Fe strain in this life-threatening infection. Briefly, upon creation of a sterile vegetation of the heart endothelium, the animals were systemically infected with an inoculum containing equal amounts of OG1RF, ΔRS05575, Δ5Fe, and Δ5FeΔRS05575 strains, and the percentage of each strain recovered from infected heart vegetations assessed 24-hours post-infection (Fig 8A). The Δ5Fe (∼12%) and Δ5FeΔRS05575 (less than 5%) strains were recovered at significantly lower rates when compared to OG1RF (∼48%) (Fig 8B). The ΔRS05575 strain was also recovered at lower rates (∼30%) but this difference was not statistically significant when compared to OG1RF.

**Fig 8.**
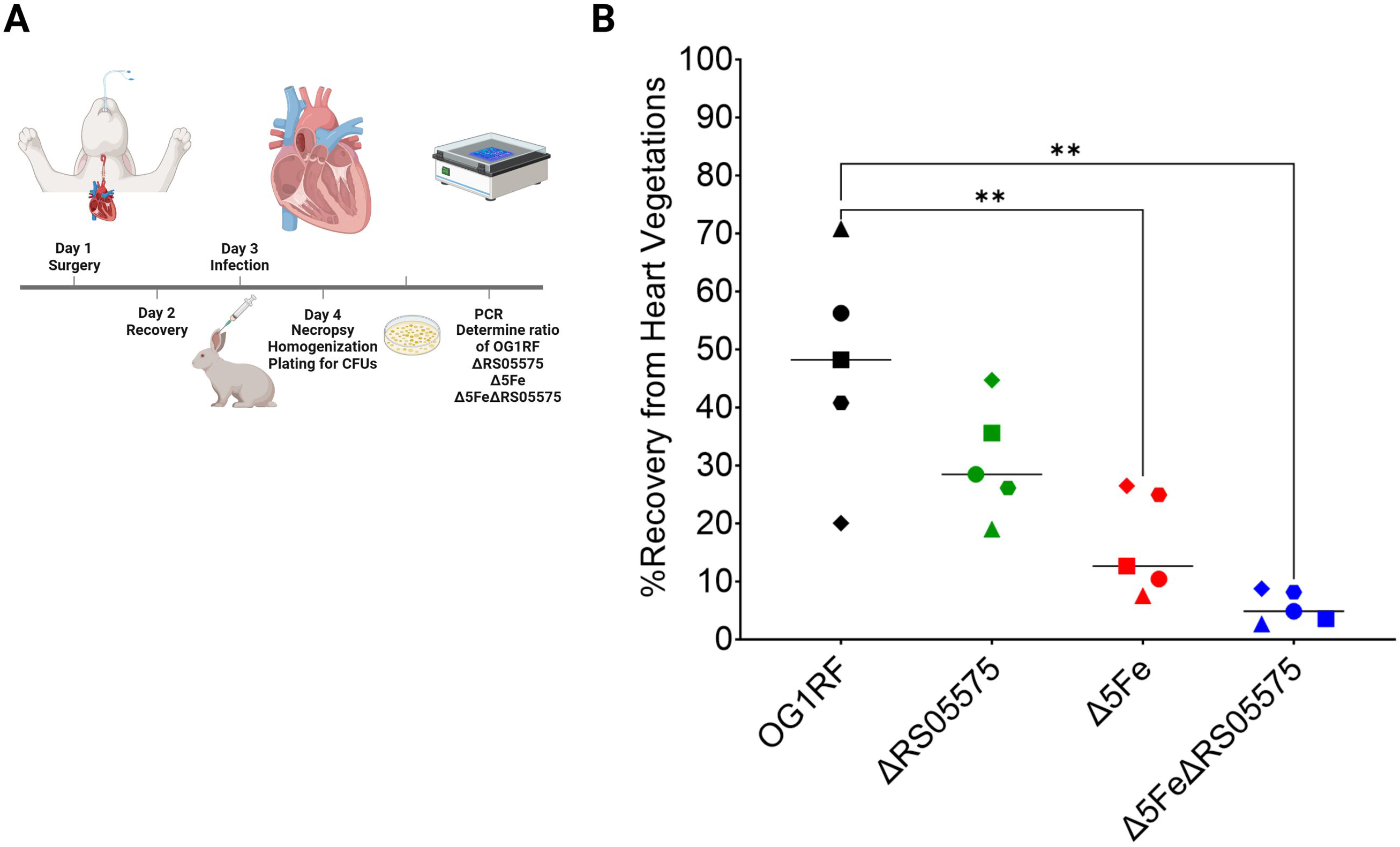
Competitive fitness of OG1RF, ΔRS05575, Δ5Fe, and Δ5FeΔRS05575 in a rabbit infective endocarditis model. Bacteria was co-inoculated (1:1:1:1 ratio) into the ear vein of rabbits 48-hours after catheter implantation. After 24-hours animals were euthanized and bacterial burdens determined in the heart vegetations. (**A**) Schematic of model. (**B**) Graph shows the percent of each strain recovered from each animal. Each symbol represents an individual rabbit, and the horizontal line represents the median recovery of each strain. Statistical significance was determined using a repeated measures one-way ANOVA with a Holm-Šidák’s multiple comparisons test, ** p≤0.01.

In the final set of experiments, we evaluated the importance of iron scavenging and RS05575 to the ability of *E. faecalis* to colonize its natural habitat, the mammalian gut. For this, we used a mouse model (32, 33) in which the gut flora is depleted with antibiotics prior to oral gavage with individual strains (Fig 9A). Strain fitness was determined by enumeration of bacteria recovered from feces 1, 2 and 3 days post-gavage (See Fig 9A and methods for details). When compared to animals infected with OG1RF, we observed significant decreases in the recovery of all three mutants over time, with the sextuple mutant showing the largest defect (Fig 9B). These studies collectively demonstrate that RS05575 enhances enterococcal fitness and virulence within the host, implicating, for the first time, anaerobic heme degradation in bacterial pathogenesis.

**Fig 9.**
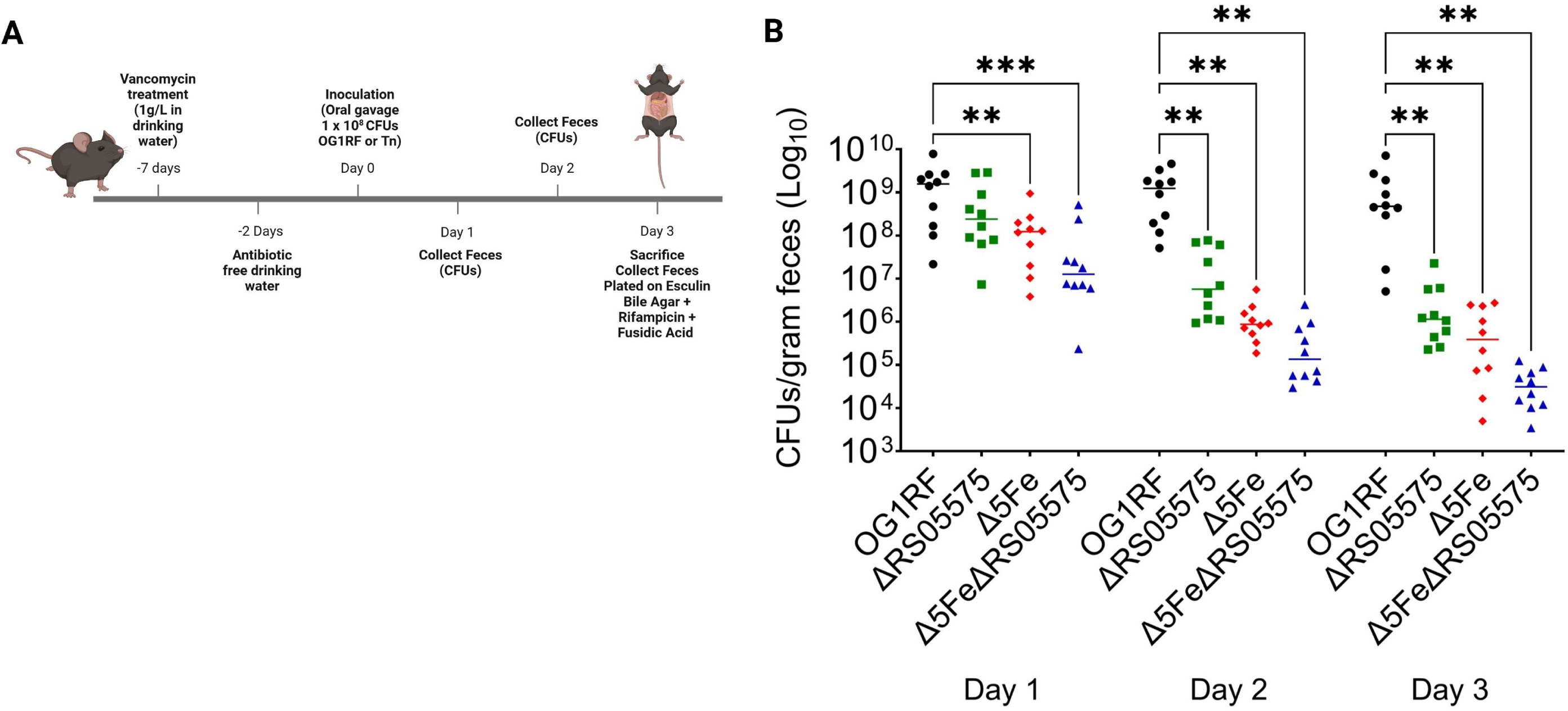
Competitive fitness of OG1RF, ΔRS05575, Δ5Fe, and Δ5FeΔRS05575 in a mouse gut colonization model. Seven-week-old C57BL6J mice were given vancomycin in drinking water to deplete endogenous enterococci prior to inoculation with *E. faecalis* strains by oral gavage. Feces were collected every day for 3 days for CFU determination. (**A**) Schematic of gut colonization model. (**B**) Colonization of mice by OG1RF, ΔRS05575, Δ5Fe, or Δ5FeΔRS05575 was determined by plating stool samples on bile esculin agar with rifampicin (200 µg mL^-1^). Data are shown with median. Statistical significance was determined by multiple Mann-Whitney tests for significance with Bonferroni-Dunn correction for multiple comparisons, *p≤0.05, **p≤0.01, ***p p≤0.001, and ****p≤0.0001.

## DISCUSSION

In bacteria, heme serves both as a nutrient cofactor and as an iron source. However, when in excess, it disrupts the cellular membrane, triggers DNA damage, and oxidizes lipids (34). To maintain heme homeostasis, bacteria evolved different mechanisms to acquire, export, synthesize, degrade, and sequester heme (13, 35). While several of these mechanisms have been identified and characterized in other Gram-positive pathogens, little is known about the mechanisms utilized by enterococci to acquire, utilize and maintain heme homeostasis (26, 34). Previously, we showed that iron starvation in *E. faecalis* can be fully reversed by heme supplementation (9). Here, we provided unequivocal evidence that heme serves as a major, if not the preferred, iron source for *E. faecalis* by showing that free iron uptake is inhibited by heme supplementation. Furthermore, we showed that intracellular heme remains elevated when free iron is abundant, indicating that *E. faecalis* possesses dedicated, likely inducible, mechanisms to degrade and then use heme as an iron source. While the prevailing bacterial mechanism to degrade heme is through oxidative degradation, mediated by heme oxygenases (17, 36), extensive bioinformatic searches for the presence of these enzymes in enterococcal genomes failed to reveal potential candidates. Thus, it is possible that enterococci rely on a non-enzymatic mechanism, termed coupled oxidation, to degrade heme when in the presence of oxygen (26, 37, 38). For example, the respiratory pathogen *Streptococcus pneumoniae* has been shown to degrade heme via production of H_2_O_2_, a metabolic byproduct of pyruvate oxidase and lactate oxidase enzymatic reactions (37–39). While *E. faecalis* does not encode either of these enzymes, it is known to generate low amounts of H_2_O_2_ that can be enhanced when cells are grown on alternative sugars such as glycerol and galactose (40, 41).

While studies to elucidate the mechanisms of aerobic heme degradation and identify the mechanism(s) by which *E. faecalis* obtains heme from the environment are active areas of investigation in our laboratory, here we described the identification of RS05575, an enzyme that resembled *E. coli* ChuW. The discovery of ChuW unveiled a new paradigm for heme degradation that, due to the high oxygen sensitivity of this new class of enzyme, is anticipated to be restricted to facultative or strict anaerobes (22, 23, 25). Despite RS05575 annotation as a coproporphyrinogen synthase, an enzyme that catalyzes the conversion of coproporphyrinogen III to protoporphyrinogen IX, *E. faecalis* genomes lack the remaining biosynthetic operon for anaerobic heme synthesis. In fact, a homologue of RS05575 in *S. aureus* Newman strain (68% amino acid similarity with RS05575) was found to have no role in anaerobic heme biosynthesis (42). Leveraging the fact that the Δ5Fe strain heavily depends on heme to maintain iron homeostasis, we showed that even though Δ5Fe strains are primed for heme uptake, the accelerated heme degradation that is observed in Δ5Fe is completely lost in the sextuple Δ5FeΔRS05575 mutant under oxygen-depleted conditions. In agreement with the anticipated oxygen-sensitivity of RS05575, we also found that RS05575 can only protect *E. faecalis* from heme toxicity under strict anaerobic conditions. Collectively, these studies reveal that RS05575 mediates heme degradation and is critical for heme homeostasis under anaerobiosis and, possibly, microaerophilic conditions.

Previously, we used the peritonitis model to demonstrate that the importance of iron scavenging transport systems to *E. faecalis* virulence was host niche dependent, and speculated that these tissue/organ-specific differences were linked to differences in heme bioavailability (9). Here, we followed up on these observations by revisiting the virulence potential of Δ5Fe in the peritonitis model while also testing the virulence potential of ΔRS05575 and Δ5FeΔRS05575. To further probe the proposed niche-specific association of iron and heme bioavailability with pathogenesis, we sampled additional organs by determining bacterial burden in livers, kidneys, and hearts homogenates. We confirmed the impaired (niche-dependent) virulence phenotype of the Δ5Fe strain that now includes evidence of impaired colonization of kidneys but no of other organs such as spleen (shown before), heart, or liver. Noteworthy, the ΔRS05575 and Δ5FeΔRS05575 strains also displayed impaired ability to colonize the kidney.

Here, it should be noted that the kidney is highly susceptible to heme-iron injury and that HO-1 levels are elevated in kidneys to protect the organ from heme toxicity (43). In the end, the most relevant finding from these studies is that virulence of ΔRS05575 alone is attenuated and exacerbated when RS05575 is inactivated in the Δ5Fe background. In fact, only the sextuple Δ5FeΔRS05575 strain displayed significant defects in dissemination to spleen and heart and was the least fit strain in the competitive rabbit IE model. Similar trends were noted in the gut colonization mouse model whereby all mutants colonized the gut poorly when compared to the parent strain. In the future, it will be interesting to assess the virulence potential of these mutants in localized infections, such as wounds or urinary tract infections, and to test their ability to colonize the gut when levels of heme are elevated, whether from intake of a heme-rich diet or due to colitis (44, 45). As we observed fitness defects in the colonization of the polymicrobial mouse gastrointestinal tract, it would also be compelling to determine how loss of RS05575, alone or in the Δ5Fe background, affects *E. faecalis* fitness and pathogenic behavior in polymicrobial biofilm infections.

While biochemical studies are still lacking, this study provides the first description of an active mechanism of anaerobic heme degradation in a Gram-positive bacterium. Moreover, it links, also for the first time, anaerobic heme degradation with bacterial colonization of the host and virulence. Because RS05575 orthologs are present in other Gram-positive bacteria, findings from this study provide the foundation for future studies that can establish a new paradigm for how other Gram-positive facultative anaerobes utilize heme and, at the same time, protect itself from heme toxicity under oxygen-depleted conditions.

## MATERIALS AND METHODS

### Bacterial strains and growth conditions

Bacterial strains used in this study are listed in Table 1. All *E. faecalis* strains were grown overnight aerobically at 37°C in BHI (Difco) unless otherwise noted. For controlled growth under metal-depleted conditions, we used the chemically defined FMC media originally developed for cultivation of oral streptococci (36), with minor modifications. The recipe for FMC is shown in Table S2. Specifically, the base media was prepared without any of the metal components (magnesium, calcium, iron, and manganese) and treated with Chelex (BioRad) to remove contaminating metals. The pH was adjusted to 7.0 and filter sterilized. All FMC component solutions were prepared using National Exposure Research Laboratory (NERL) trace metal grade water, filter sterilized, and then added to the media. Heme (Sigma-Aldrich) was prepared in 1.4 M NaOH in NERL trace metal grade water. Calcium, magnesium, manganese, iron, and heme were added at concentrations specified in the text or figure legend. For reverse transcriptase quantitative PCR (RT-qPCR) analysis, RNA was isolated from cells grown in FMC[-Fe] to OD_600_ of 0.4 and spiked with 20 µM heme with aliquots taken 0 and 60 minutes post-heme supplementation. To generate growth curves, cultures were grown in FMC[-Fe] and diluted 1:200 into fresh FMC[-Fe] supplemented with heme and/or FeSO_4_ as indicated in the text and figure legends. Aerobic cell growth was monitored using the Bioscreen growth reader (Oy Growth Curves). Growth of anaerobically grown cells was monitored in a 96-well plate reader (Byonoy) in an anaerobic chamber (Coy).

**Table 1.** Bacterial strains used in this study.

| Strains | Relevant Characteristics | Source |
| --- | --- | --- |
| <u>*E. faecalis*</u> |  |  |
| OG1RF | Rif<sup>R</sup> Fus<sup>R</sup> | Lab collection |
| $\Delta efaCBA\Delta feoB\Delta fhuB\Delta fitAB\Delta emtB$<br>($\Delta 5Fe$) | *efaCBA* deletion; *feoB* deletion; *fhuB* deletion; *fitAB* deletion; *emtB* deletion. | (9) |
| $\Delta RS05575$ | RS05575 deletion. | This study |
| $\Delta RS05575c$ | RS05575 complemented with reintegration. | This study |
| $\Delta 5Fe\Delta RS05575$ | RS05575 deletion; *efaCBA* deletion; *feoB* deletion; *fhuB* deletion; *fitAB* deletion; *emtB* deletion. | This study |
| $\Delta 5Fe\Delta RS05575c$ | RS05575 deletion; *efaCBA* deletion; *feoB* deletion; *fhuB* deletion; *fitAB* deletion; *emtB* deletion. RS05575 complemented with reintegration. | This study |
| CK111 | OG1S *upp4::P23repA4*, Spec<sup>R</sup>.<br>Conjugation donor strain. | (46) |

### Construction of mutant strains

Markerless deletions of RS05575 in *E. faecalis* OG1RF were carried out using the pCJK47 genetic exchange system (46). Briefly, PCR products with ∼1 kb in size flanking each coding sequence were amplified with the primers listed in Table S3. To avoid unanticipated polar effects, amplicons included either the first or last residues of the coding sequences. Cloning of amplicons into the pCJK47 vector, electroporation, and conjugation into *E. faecalis* strains and isolation of single mutant strains (ΔRS05575) were carried out as previously described (46). Isolation of Δ5FeΔRS05575 was done by conjugation of the pCJK47 vector with the Δ5Fe strain used in a previous publication (9). All gene deletions were confirmed by PCR sequencing of the insertion site and flanking region.

### Construction of the complemented strains

The pCJK47 vector was used to insert RS05575 back into its original genetic loci to be regulated by the native promoter. Briefly, the coding sequence of RS05575 was amplified from OG1RF using the primers listed in Table S3. We used the In-fusion cloning system (Takara Bio) to generate the allelic exchange plasmid. The pCJK47 vector was digested with BamHI and PstI to yield pCJK47-RS05575c vector. Upon propagation in *E. coli* EC1000, pCJK47-RS05575c was electroporated into the conjugation strain *E. faecalis* CK111, and the plasmid mobilized into ΔRS05575 using a standard conjugation protocol (46).

### RNA analysis

RNA was isolated from cells before and after exposure to 20 µM heme using the PureLink™ RNA Mini Kit (Invitrogen). Genomic DNA (gDNA) was degraded using TURBO Dnase kit (Invitrogen) and cDNA synthesis from 1 µg total RNA using the high-capacity cDNA reverse transcription kit (Applied Biosystems). RT-qPCR was performed using iTaq Universal SYBR supermix (BioRad) with primers listed in Table S3. Copy number was determined using a standard curve generated from OG1RF genomic DNA (gDNA) and fold change calculated.

### Phylogenetic analysis

ChuW and RS05575 amino acid sequences were compared using Clustal-Omega multiple sequence alignment and Needleman-Wunsch global alignment tools on SnapGene. BlastP searches in both NCBI and BV-BRC databases were used to identify homologues of RS05575 in other bacteria. Select homologues were used to generate a multiple sequence alignment for phylogenetic tree using EMBL-EBI’s Clustal-Omega. The phylogenetic trees were then modified for readability using the interactive Tree of Life (iTOL) version 6.

### Protein structure predictions

Tertiary structures of *E. faecalis* RS05575 and *E. coli* ChuW were obtained using AlphaFold2 Colab notebook and a predictive structure of *E. faecalis* RS05575 binding heme was generated using AlphaFold server. All image files (PDB) were constructed using ChimeraX1.3 (47–49). Structural alignments were performed on the Research Collaboratory for Structural Bioinformatics website using AF-A0A0M2ASD4-F1 (RS05575) and AF_AQFA0A384LP51F1 (ChuW) (31).

### ^55^Fe uptake

Overnight cultures of OG1RF were grown in FMC[-Fe]. Cultures were grown to mid-log phase (OD_600_ ∼0.5), at which point 1 µM ^55^Fe (Perkin-Elmer), with and without competing cold metals, was added to each culture followed by incubation at 37°C. Immediately after ^55^Fe addition and 1 and 5 minutes after, 200 µL aliquots were transferred to a nitrocellulose membrane pre-soaked in 1 M NiSO_4_ solution (to prevent nonspecific binding) and placed in a slot blot apparatus. Free ^55^Fe was removed by four washes in 100 mM sodium citrate buffer using vacuum filtration. The membranes were air dried, cut, and dissolved in 4 mL scintillation counter cocktail. Radioactivity was measured by scintillation with “wide open” window setting using a Beckmann LSC6000 scintillation counter. The count per million (cpm) values from ^55^Fe free cells were obtained and subtracted from the cpm of treated cells. The efficiency of the machine was ∼30.8% and was used to convert cpm to disintegrations per minute (dpm), which was then converted to molarity and normalized to CFU.

### Intracellular heme quantification

Overnight cultures were grown in FMC[-Fe] under aerobic conditions, and sub-cultures grown under varying degrees of oxygen to OD_600_ 0.4 in FMC +/- iron and/or +/- heme. Specifically, cultures were grown with 50% headspace in a shaking incubator, statically with 50% headspace in an aerobic incubator, statically with no headspace in an aerobic incubator, or statically with no headspace in an anaerobe chamber using media with dissolved oxygen (designated as FMC_DO_ in the text). For heme uptake and degradation kinetic experiments, cultures were first grown in FMC[-Fe] lacking dissolved oxygen (designated as FMC_O2P_[-Fe]) to OD_600_ 0.4, 10 µM heme was then spiked into the cultures and aliquots taken after 5, 15, and 60 minutes. Degradation of intracellular heme was assessed by growing cells in FMC_O2P_ [-Fe] to OD_600_ 0.4, spiking cultures with 10 µM heme for 15 minutes, and then washing cultures in 0.5mM EDTA in NERL grade metal free PBS once, and in NERL grade metal free PBS twice. The cultures were then resuspended in FMC_O2P_[-Fe] and samples taken at 0, 60, and 180 minutes after removal of heme. Cells were washed at least 3 times in 1X PBS and lysed using bead beating in 1 mL NERL trace metal grade water. Lysates were used to determine heme content using a heme detection kit (Sigma-Aldrich) and normalized to protein content using the BCA assay (Sigma-Aldrich).

### Intraperitoneal challenge mouse model

The model has been described previously (43) such that only a brief overview is provided below. To prepare the bacterial inoculum, bacteria were grown in BHI to an OD_600_ of 0.5, the cell pellets collected, washed once in 0.5 mM EDTA and twice in trace metal grade PBS, and suspended in PBS at ∼2 x 10^8^ CFU mL^-1^. Seven-week-old C57BL6J mice purchased from Jackson laboratories were intraperitoneally injected with 1 mL of bacterial suspension and euthanized by CO_2_ asphyxiation 48-h post-infection. The abdomen was opened to expose the peritoneal lining, 5 mL of cold PBS injected into the peritoneal cavity with 4 mL retrieved as the peritoneal wash content. Quantification of bacteria within the peritoneal cavity was determined by plating serial dilutions on tryptic-soy agar (TSA) containing 200 µg mL ^-1^ rifampicin and 10 µg mL^-1^ fusidic acid. For bacterial enumeration inside spleens, livers, kidneys, and hearts, organs were surgically removed, rinsed in 70% ethanol to remove bacteria attached to the exterior of the organ, rinsed in sterile PBS, homogenized in 1 mL PBS, serially diluted, and plated on selective TSA plates. These experiments were approved by the University of Florida Institutional Animal Care and Use Committee (protocol 202200000241).

### Infective endocarditis rabbit model

Pathogen-free New Zealand White rabbits (2-4kg; Charles River) were utilized in an endocarditis model as described previously (29). Prior to surgery, rabbits were anesthetized with ketamine, xylazine, glycopyrrolate, buprenorphine, isoflurane, and sevoflurane, with bupivacaine applied locally. A PE-90 catheter (Becton-Dickinson) was inserted into the aortic valve via the right carotid artery; placement was confirmed by ultrasound. Each catheter was tied off and sutured in place, and the incision was closed with staples. Rabbits were monitored for the next 48 hours to ensure stability prior to infection. Bacterial inoculum was prepared by growing cells in BHI. Each strain was washed as described above and normalized to OD_600_ ∼0.8 in Chelex-treated (BioRad) PBS. An inoculum was prepared by combining equal volumes of each strain, achieving a total inoculum of 6 x 10^7^ CFUs mL^-1^; 0.5 mL was then delivered via ear vein injection. From the inoculum, 1 mL was plated and used to verify equal distribution of each strain by PCR using primers listed in Table S3. 24 hours post infection, rabbits were sedated by intramuscular injection with acepromazine (Covetrus) and then euthanized via ear vein injection of Euthasol (Med-Pharmex). Harvested vegetations were placed into PBS, homogenized, and plated on BHI agar. At least 250 colonies per rabbit were analyzed by PCR to determine the percent recovery of each strain that was determined by dividing the number of each specific strain recovered by the total number of colonies assayed and then multiplying by 100. These experiments were approved by the Virginia Commonwealth University Institutional Animal Care and Use Committee (protocol AM10030).

### Intestinal colonization mouse model

Seven-week-old C57BL6 male mice were purchased from Jackson Laboratories and given one week to equilibrate their microbiota prior to experimentation. Mice were given antibiotics (0.5 mg/mL cefoperazone + 1 mg/mL vancomycin) in drinking water *ad libitum* for 5 days followed by a 2-day recovery period and subsequent infection. Mice were confirmed culture-negative for endogenous enterococci after vancomycin treatment via selective plating as described below. Mice were infected via oral gavage with 5 × 10^8^ CFUs of *E. faecalis* dissolved in PBS. Enterococcal CFUs were quantified daily from fecal samples. Samples were diluted and homogenized in PBS and serially plated onto bile esculin agar for total enterococci. To distinguish *E. faecalis* lab strains from endogenous enterococci, samples were also grown on bile esculin agar with rifampicin (200 µg/mL). These experiments were approved by the Animal Care and Use Committees of the Children’s Hospital of Philadelphia (protocol IAC 21–001316).

### Statistical analyses

All data sets were analyzed using GraphPad Prism 10 software. Statistical significance in the transcriptional expression studies were analyzed by comparing the fold change in copy number before and after heme supplementation using a Student’s T-test.

Statistical differences in ^55^Fe uptake were determined by Two-way ANOVA and Dunnett’s multiple comparison test. Intracellular heme content of OG1RF grown in the presence or absence of oxygen, with or without an excess iron source was analyzed by Student’s T-test. The intracellular heme content of strains grown in decreasing oxygen levels was analyzed with a Two-way ANOVA and Šidák’s multiple comparison test. Statistical significance in the mouse peritonitis and in the gut colonization model were determined by the Mann Whitney test.

Statistical differences in recovery of strains from the competitive rabbit IE model was determined by a repeated measure One-way ANOVA with a pairwise Holm Šidák’s multiple comparison test.

## ACKNOWLEDGEMENTS

We thank Drs. Jennifer Bradley, Liang Bao, and Josephina Vossen and Ms. Kali Williams, Katherine Atran, Valerie Assi and Nicai Zollar for assistance with the rabbit endocarditis model. This study was supported by NIH-NIAID grant R21 AI137446 to J.A.L and R35GM138369 to J.P.Z. D.N.B. was supported by NIH-NIDCR training grant T90 DE021990 and by American Heart Association predoctoral fellowship 907592. J.P.Z. was also supported by the Center for Microbial Medicine at the Children’s Hospital of Philadelphia.

